# Aerobic Exercise Intensity: A Dose-Response Effect on Motor Adaptation and Learning

**DOI:** 10.1101/2025.10.28.680122

**Authors:** Nesrine Harroum, Yasmine Mahrez, Benjamin Pageaux, Jason L. Neva

**Affiliations:** École de kinésiologie et des sciences de l’activité physique (EKSAP), Faculté de médecine, Université de Montréal, Montreal, QC, H3C 3J7, Canada; Centre de recherche de l’Institut universitaire de gériatrie de Montréal (CRIUGM), Montreal, QC, H3W 1W4, Canada; Centre interdisciplinaire de recherche sur le cerveau et l’apprentissage (CIRCA), Montreal, QC, H3C 3J7, Canada

**Keywords:** acute exercise, exercise intensity, visuomotor adaptation, visuomotor rotation task

## Abstract

Acute aerobic exercise (AEX) can enhance motor learning. While AEX intensity likely plays a key role, there is mixed evidence for AEX-enhanced motor skill acquisition and learning across a spectrum of exercise intensities. This may stem, in part, from inconsistent AEX parameters (i.e., intensity, structure, and duration) employed within and across studies. Additionally, evidence suggests that AEX can enhance a specific form of motor learning, namely motor adaptation. Moderate- and high-intensity AEX can increase motor adaptation, but evidence remains limited and inconsistent. Hence, the impact of AEX intensity on motor adaptation remains unclear. Here, we investigated the influence of AEX intensity on motor adaptation, while controlling for AEX structure and duration. Eighty young adults were assigned to four cycling AEX/Rest groups (n=20/group): 20 min of light (LIIT), moderate (MIIT), or high (HIIT) intensity interval training, or Rest (control). AEX consisted of four 3-min cycling intervals (LIIT, 35% heart rate reserve [HRR]; MIIT, 55%HRR; HIIT, 80%HRR) and 2-min active recovery (25%HRR). Participants practiced a visuomotor rotation task immediately after AEX/Rest (adaptation) and at a no-AEX 24 h retention test (motor learning). We found that: (1) all AEX intensities enhanced motor learning compared to Rest, and (2) HIIT enhanced motor adaptation and learning to the greatest extent, followed by MIIT then LIIT. This is the first study to demonstrate a dose-response effect of AEX intensity on motor adaptation and learning. Our results highlight the importance of considering intensity when prescribing AEX in sports and clinical contexts to promote motor learning.

## INTRODUCTION

A growing body of evidence demonstrates that acute aerobic exercise (AEX) enhances motor learning^1–5^. Specifically, AEX performed before motor task practice can enhance skill acquisition^5–8^ and motor learning as demonstrated by sustained performance improvement assessed at delayed retention tests 5 h^9^ to 7 days later^3–5,10,11^. Importantly, a previous meta-analysis demonstrated that exercise parameters (e.g., intensity) play a critical role in the impact of AEX on motor learning^1^.

The beneficial impact of AEX on motor learning has been demonstrated across a spectrum of AEX intensities, including light, moderate and high^2–4,10–14^. A recent meta-analysis^1^ further refined this understanding, suggesting that moderate-intensity AEX tends to enhance motor skill acquisition (i.e., immediate post-exercise performance^5–8^), whereas higher intensity AEX more robustly enhances motor learning, as assess during delayed retention tests (i.e., 24 h after practice^1–4,10–13^). Despite these insights, findings are variable. This variability may stem, in part, from inconsistent AEX parameters (i.e., intensity, structure and duration) employed both within and across studies^6,14–16^. Additionally, most research has evaluated the impact of a single AEX intensity compared to rest^2,3,5,7,9–12,17,18^. Few studies have directly compared the effects of two different AEX intensities, and those that have often varied exercise structure or duration (e.g., high-intensity *interval* vs. moderate-intensity *continuous*^6,14–16^) or kept structure constant (e.g., moderate-intensity *continuous* vs. light-intensity *continuous* AEX^13,19–21^). Given this inconsistent employment of AEX intensity, structure and duration, the observed variability in motor learning outcomes is unsurprising. Indeed, some studies report greater enhancement from light-intensity^13,20^, or moderate-intensity^6,16^, compared to high-intensity AEX. Conversely, others found equivalent impact following moderate- and high-intensity AEX (20 min interval^19^), as well as light- and high-intensity (19 min interval^22^), even when similar structures and durations were employed (28 or 15 min continuous^15^; 25 min continuous and 23 min interval^14^). Thus, the precise impact of AEX intensity on motor learning remains unclear. Crucially, no study to date has examined the dose-response effect of AEX across light, moderate and high intensities while rigorously controlling for exercise structure and duration.

Another important factor to consider in the impact of AEX on motor learning is the motor task type investigated^1^. Most previous studies have used visuomotor tracking and sequence tasks, with few investigating the impact of AEX on motor adaptation. Motor adaptation is identified as a specific category of motor learning^23,24^ that is distinct from other forms and is associated with unique neurophysiological and behavioral characteristics^16,17–20,25–29^. Recent work shows that AEX can enhance motor adaptation and learning^1,5,17,30^, yet not all studies show these effects^30^. Specifically, a single bout of moderate-intensity *continuous*^5^ or high-intensity *interval*^17^ AEX can enhance adaptation and learning of a visuomotor rotation task. High-intensity AEX can enhance performance at a visuomotor rotation task 1 h following practice^17^ and moderate-intensity^5^ AEX can enhance performance immediately and 24 h following initial practice, suggesting AEX at both these intensities enhances motor adaptation and learning. In contrast, studies have found that moderate-^30^ or high-intensity^17^ AEX does not enhance motor adaptation immediately following exercise, and high-intensity AEX does not enhance motor adaptation assessed at 24-h and 7-day retention tests^17^. Further, while there are few studies that have investigated the effects of light-intensity AEX on motor learning^13–15,21,22^, no study to date has examined its effects on motor adaptation. Taken together, due to the limited number of studies, which used various AEX parameters, the critical role of AEX intensity to enhance motor adaptation and learning remains unclear.

This study aimed to determine the dose-response effect of AEX across the full physiological continuum of intensity (light, moderate, high) on motor adaptation and learning, while controlling AEX duration and structure. Consistent with previous literature showing enhanced motor learning following moderate- and high-intensity exercise^1,24,31^, we hypothesized that moderate- and high-intensity AEX would lead to superior motor adaptation and learning compared to rest.

## METHODS

### Participants

Characteristics of participants are presented in *Table 1*. All data are presented as mean ± standard deviation (SD) unless otherwise noted. Eighty healthy young adults (25 ± 5 years old, 50% F) took part in the study. Using G*Power, a sensitivity analysis with an alpha risk of 0.05, a total sample size of 80 with 4 groups and 25 measurements (blocks), revealed that we had a 80% chance of observing a small effect size f(U) = 0.14 (∼ *η²_p_* = 0.02) or higher, for repeated measures ANOVA within-between interaction^32^. Participants reported being free of neurological disorders, psychiatric disorders, musculoskeletal injuries or chronic illnesses (e.g., cardiovascular disease) and no vision impairment. The Ethics Committee of the Centre de recherche de l’Institut Universitaire de Gériatrie de Montréal approved all experimental procedures (CER VN 23-24-24).

**Table 1.**
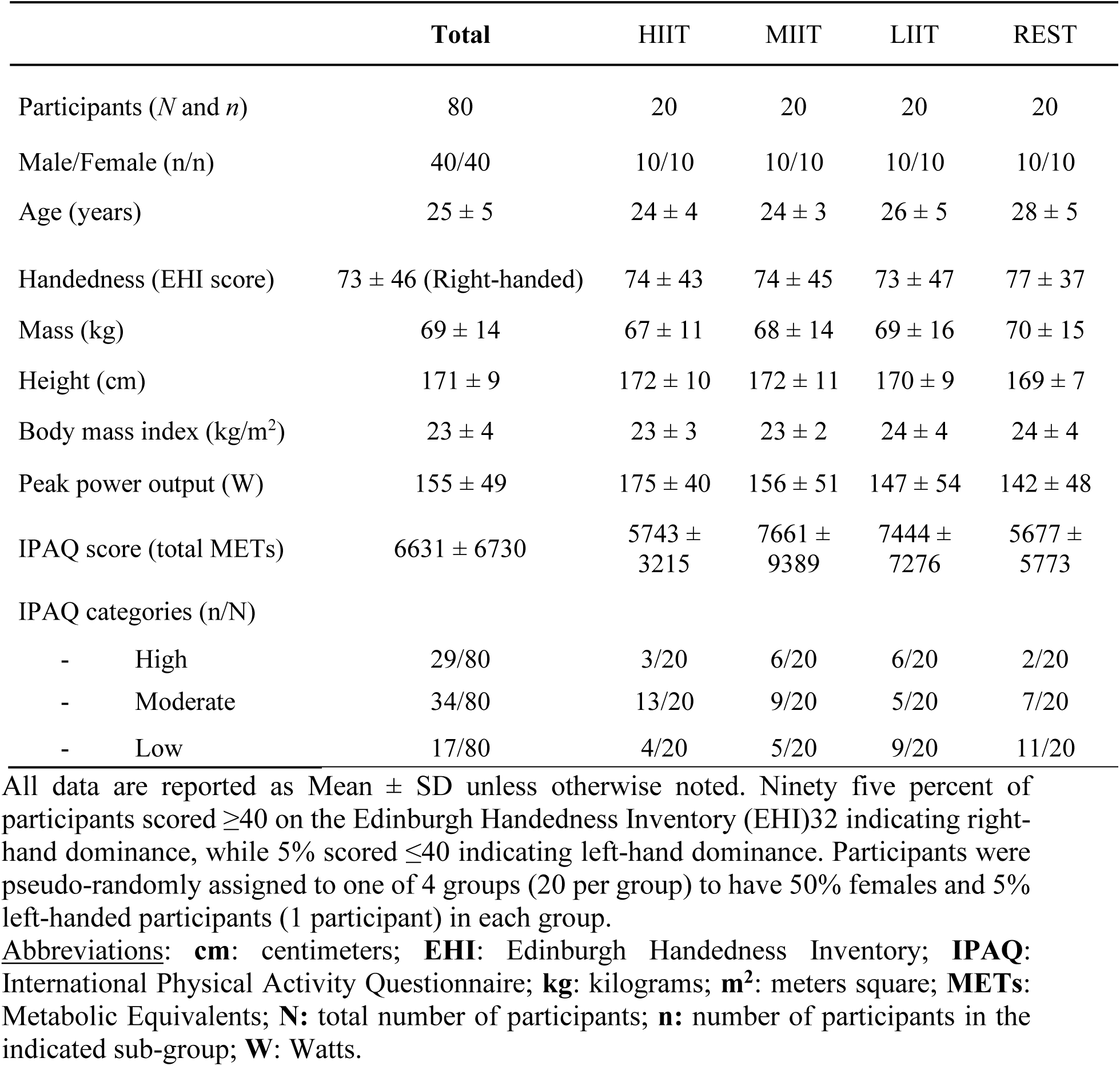
Participants characteristics.

### Experimental design

All participants completed a preliminary visit and 2 experimental sessions with a minimum of three days between the preliminary and the first experimental session, and 24 h between the first and second experimental sessions (Fig. 1). The experimental sessions aimed to assess the impact of AEX intensity on *i)* motor adaptation (visuomotor rotation task performance immediately following exercise) and *ii)* motor learning (visuomotor rotation task performance at the 24 h retention test). During the preliminary visit, participants completed questionnaires on hand dominance using the Edinburgh Handedness Inventory^33^ and daily physical activity using the International Physical Activity Questionnaire (IPAQ)^34^. Then, following 5 min at rest to obtain resting heart rate (*HRrest*), participants underwent an incremental exercise test to exhaustion to determine peak power output and peak heart rate (*HRpeak*). Resting heart rate and peak heart rate were used to prescribe the exercise intensity during the subsequent sessions based on heart rate reserve, and peak power output was used for categorization of intensity during the AEX sessions. Following the preliminary visit, participants were pseudo-randomly assigned to one of the following four groups: 20 minutes of *i)* seated rest, *ii)* light-intensity interval training (LIIT), *iii)* moderate-intensity interval training (MIIT), or *iv)* high-intensity interval training (HIIT) AEX. During the first experimental session, immediately following AEX or rest, participants practiced a visuomotor rotation task^17,35–37^. Visuomotor rotation task practice during this experimental session served to assess motor adaptation. Participants returned 24 h later to perform a no-AEX/Rest retention test for the second experimental session, which assessed motor learning^5,17^. All experimental sessions to assess motor adaptation and learning were performed at the same time of day (± 3 h)^38^.

**Fig.1.**
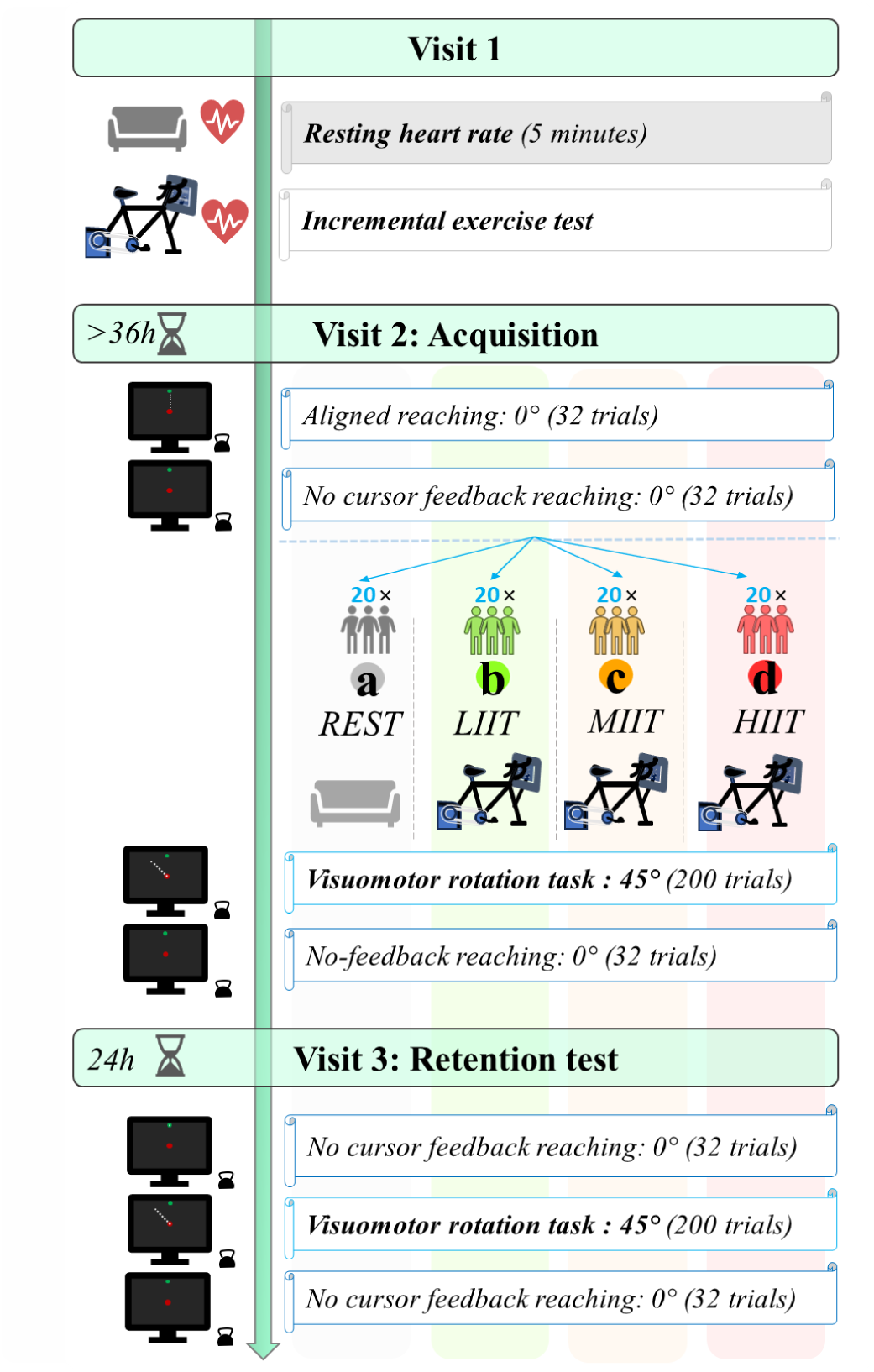
Study Design: An incremental exercise test to exhaustion was conducted in visit 1 (preliminary visit) to establish the exercise intensities for the four subsequent visits (experimental sessions) following ACSM recommendations based on the heart rate reserve. The four experimental sessions were conducted on visit 2 and involved 20 minutes of either (a) seated rest, (b) light-intensity interval cycling (LIIT), (c) moderate-intensity interval cycling (MIIT), or (d) high-intensity interval cycling (HIIT). Participants were randomly assigned to each group. Immediately (Visit 2) and 24h (Visit 3) after each experimental session, participants performed a visuomotor rotation task. <u>Abbreviations</u>: ***°***: degrees; **AEX**: acute aerobic exercise (4 blocks of 3 minutes cycling followed by 2 minutes active recovery at 25% heart rate reserve); **h**: hours; **HIIT**: high interval training cycling acute exercise (80% heart rate reserve); **LIIT**: light interval training cycling acute exercise (35% heart rate reserve); **MIIT**: moderate interval training cycling acute exercise (55% heart rate reserve).

#### Incremental exercise test and physical activity

During the preliminary visit, participants completed an incremental exercise test on an upright cycle ergometer (Cyclus2, CY00100) to determine the group-assigned AEX intensity^39,40^. Cycling began at a power output equivalent to each individual participant’s body mass (e.g., 75W for 75kg), with intensity increasing every 2 minutes by 15, 20, 25 or 30 W, in accordance with their body mass^39,40^. Participants maintained pedaling cadence between 60 and 80 rotations per minute (rpm) until exhaustion, defined as being unable to maintain a cadence greater than 60 rpm for 10 s despite verbal encouragement. Within the last 20 seconds of each stage, participants reported their perceptual responses, which included: perception of effort intensity using the CR100 scale^41^, muscle (thigh) pain using a numerical rating scale ranging from 0 (no pain) to 10 (maximum pain)^42^ and affective response using the Feeling scale^43^. Daily levels of physical activity over the 7 days preceding the preliminary visit were assessed using the IPAQ^34^ (see Supplementary information “IPAQ description” for more details).

#### Heart rate (HR) measurement

Heart rate (HR) was monitored using a chest belt (Cyclus2, Germany) throughout the incremental exercise test and during rest. We used the percentage of heart rate reserve (%HRR) method to prescribe the cycling exercise intensities^44^. HRR refers to the difference between HR_peak_ recorded during an incremental test and the HR_rest_ measured during 5 minutes of seated rest before the incremental test. To prescribe exercise intensity, we used the Karvonen formula^45,46^:

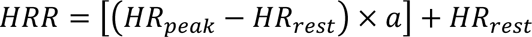

The intensities were prescribed via the constant *a* with the following values corresponding to the exercise intensities as described in the next section: 0.25, 0.35, 0.55, 0.80.

#### Acute Exercise Session

Following a 5-minute warm-up at 25% HRR on the same cycle ergometer used in the incremental exercise test, participants engaged in 20 min of interval cycling, involving four 5-minute blocks. Each block involved 3 min of pedaling at one of the target intensities: i) light (35% HRR, LIIT), *ii)* moderate (55% HRR, MIIT), or *iii)* high (80% HRR, HIIT). Each 3 min block at the target intensity was followed by 2 min of active rest (25% HRR) for all groups. Throughout the warm-up and AEX sessions, participants maintained a cycling cadence of 60-80 rpm, and reported their perceptual responses^41^ ^42^ ^43^. Specifically, these perceptual responses were reported by the participants at two time points during each block: *i)* at 0-20 seconds, and *ii)* at 2 minutes 30 seconds, from the beginning of each block. HR was continuously monitored using a heart rate belt (Polar T31C heart rate sensor). At the beginning and end of each cycling exercise interval and the warm-up, the experimenters recorded heart rate. Throughout cycling exercise, participants were instructed to maintain their hands in a relaxed position, resting them on top of the handlebars and to avoid gripping them.

#### Rest

The rest condition lasted 25 minutes to match the duration performed by the AEX groups. Participants were seated comfortably on a chair and watched an emotionally neutral documentary^47^, while HR and perceptual responses were collected as the AEX groups.

#### Visuomotor reaching task

Participants sat comfortably 1 meter in front of a 32-inch computer monitor, with their dominant hand resting on a table. Participants performed out-and-back reaching movements toward visual targets displayed on the monitor using a custom mouse (Fig. 2A). During all reaching tasks, participants were unable to see their arms or hands. Real-time hand position feedback was displayed as a white 1-cm diameter cursor on the screen. All reaching tasks involved participants moving the cursor from the central start location to a peripheral target and then reaching back to the central target after it reappeared. Cursor feedback of hand position was provided during the reach out to peripheral targets (except for no-cursor feedback reaching) and not provided on the reach back to center until the cursor was within a 3-cm radius of the central target. The central target was represented by a red 2-cm diameter circle and the eight peripheral targets were represented by 2-cm diameter green circles. The peripheral targets were radially spaced 45° apart and 15 cm from the central start target. The peripheral targets were presented randomly within blocks of 8 trials, ensuring none of the 8 peripheral targets were repeated consecutively within each block (Fig. 2A). A trial was considered successful when the participant moved the cursor from the central target to the peripheral target and remain there for 500 ms, after which the central target reappeared, and they were required to reach back to the central target. If the peripheral target was not reached within 5 seconds, the trial was considered unsuccessful. For all reaching tasks, participants were instructed to reach as quickly and accurately (i.e., the most direct path) as possible, from the central to the peripheral targets.

**Fig.2.**
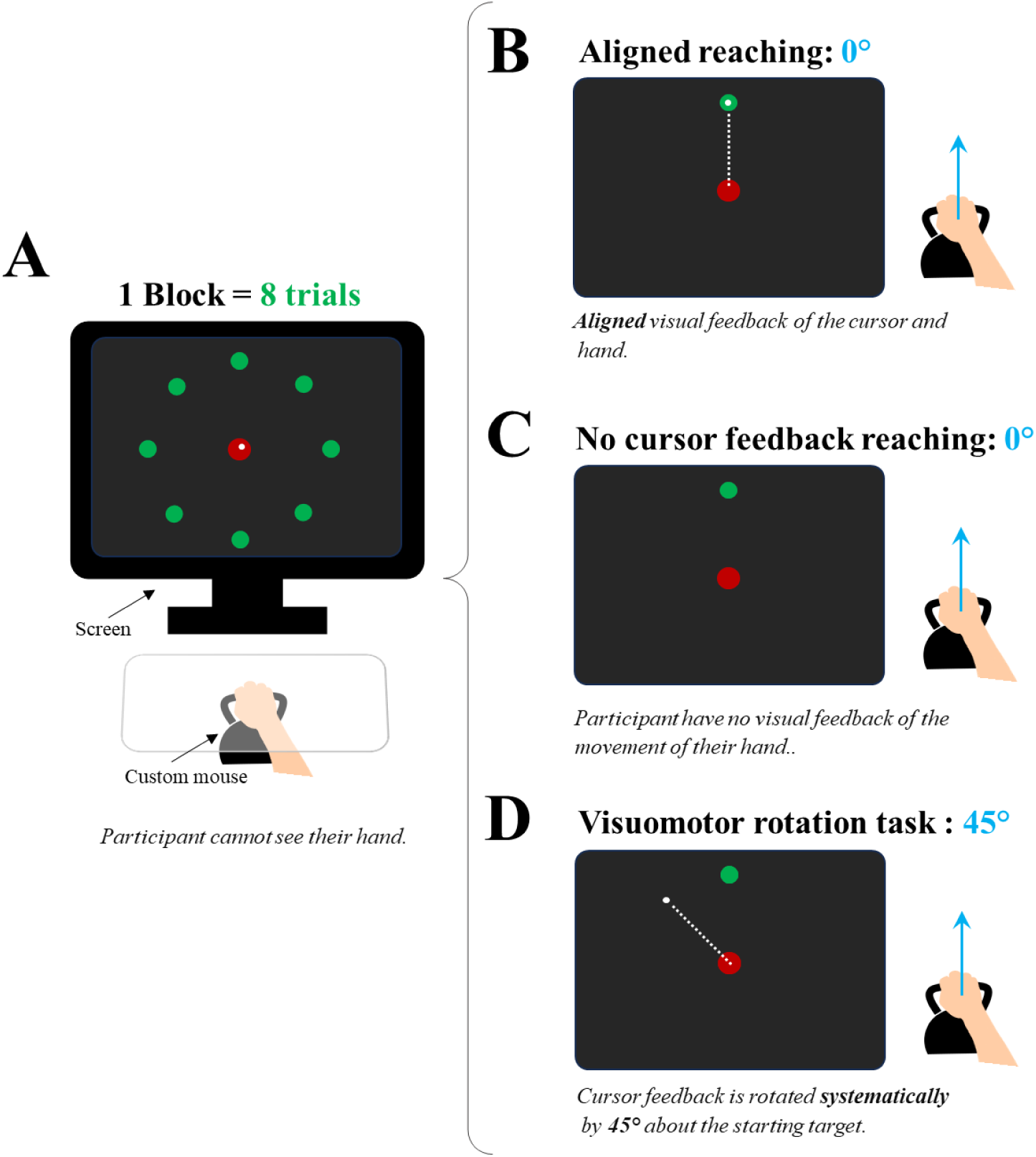
Visuomotor rotation task: **(A)** The participant sat 1 meter away from a 32-inch screen, holding a custom mouse in their hidden dominant hand. A 1 cm white cursor displayed real-time hand movements. Each block featured eight peripheral targets, spaced 45° apart and 15 cm from the center, appearing sequentially in random order without repetition. The participant moved the cursor from a 3 cm red start circle to a 2 cm green target circle as quickly and accurately as possible. **(B)** The cursor was moving (the white dotted line represents the cursor trajectory) along with the hand trajectory (blue arrow) while the participants were trying to reach the peripheral green targets. **(C)** No cursor was displayed while participants reached for the peripheral green targets. **(D)** The cursor was rotated 45° counterclockwise relative to the actual hand trajectory while reaching for the peripheral green targets. <u>Abbreviations</u>: ***°***: degrees.

Participants performed 3 reaching tasks: (1) aligned feedback reaching (Fig. 2B), (2) no-cursor feedback reaching (Fig. 2C), and (3) the visuomotor rotation task (Fig. 2D)*. Aligned feedback reaching.* Immediately before exercise or rest, participants performed reaching movements with the cursor aligned with hand movement for 32 trials (Fig. 2B). These trials served as familiarization with the general visuomotor reaching task parameters and as a baseline assessment of reaching movements without alterations of visual feedback^35,48^.

##### No-cursor feedback reaching

See supplementary Fig. 3 and supplementary Table 5 for details on the experimental procedure, statistical analysis and results.

##### Visuomotor rotation task

Immediately after AEX/Rest and at the 24-h retention test, participants practiced a visuomotor rotation task. In this reaching task, a 45° clockwise rotation was applied to the cursor’s movement relative to the actual hand trajectory about the central target^35,49^ (Fig. 2D). The visuomotor rotation task consisted of 200 trials, divided into 5 sets of 40 trials. Participants had a 30-s break following each set.

### Data processing and statistical analysis

#### Data processing

All trials for each reaching task were screened with respect to their velocity profile, movement trajectory and end-point position using custom MATLAB scripts (MATLAB R2022a, MathWorks, Natick, MA, USA). Trials with reaction time, movement time and angle at peak velocity exceeding 3 standard deviations from the mean were discarded (accounting for 0.9% of all trials). The outward reach was analyzed from the central to peripheral targets with respect to the cursor position. The return movement to the central start target was not analyzed. Reaching performance was assessed using two temporal measures and one kinematic measure. The temporal measures included *i)* reaction time, representing time (in seconds) between target appearance and initial cursor movement from the center target, and *ii)* movement time, representing the time (in seconds) from cursor movement initiation to peripheral target acquisition. The kinematic measure of interest was angle at peak velocity, representing the angular error (in degrees) at the fastest point of movement^5,50,51^.

#### Statistical analysis

Analysis of variance (ANOVA) was used to analyze visuomotor reaching task performance and the data collected during AEX/Rest (HR, perceptual responses). Details of each specific analysis are detailed below. Post hoc analyses were conducted using Holm-Bonferroni correction where appropriate. We assessed normality and homoscedasticity through residual statistics, skewness, kurtosis values, and visual inspection of plots. All statistical procedures were performed using Jamovi software (version 2.2.5), with statistical significance set at *p* < .05. Effect sizes were calculated as partial eta squared (*η²_p_*) and interpreted according to established guidelines^52^: 0.01 for small, 0.06 for moderate, and 0.14 for large effects.

##### Aerobic exercise / Rest data

To verify the impact of different AEX intensities (HIIT, MIIT, LIIT), we performed one-way ANOVAs with between-subjects factor GROUP (HIIT, MIIT, LIIT, REST) on the physiological and perceptual responses. To assess physical activity levels between groups, we performed a one-way ANOVA with a between-subjects factor of GROUP (HIIT, MIIT, LIIT, REST) on the METS calculated from the IPAQ.

### Visuomotor reaching task data

#### Aligned feedback reaching

We performed a one-way ANOVA with between-subjects factor GROUP (HIIT, MIIT, LIIT, REST) on aligned feedback reaching to characterize general visuomotor reaching task performance.

#### Visuomotor rotation task

Two-way mixed-model ANOVAs with between-subjects factor GROUP (HIIT, MIIT, LIIT, REST) and within-subject factor BLOCK (1 to 25) were conducted for each dependent measure (reaction time, movement time, angle at peak velocity). These analyses were performed separately for *adaptation* (motor practice immediately after AEX/Rest) and the *retention test* (motor practice 24 h after AEX/Rest). For *adaptation*, following significant interaction, to reduce the number of post hoc analyses, follow-up pairwise comparisons were made between groups with the first and final three blocks of visuomotor rotation task practice. Further, for *adaptation*, to ascertain whether a significant main effect of GROUP was attributable to differences in immediate post-AEX/Rest performance or emerged with continued practice, an additional a one-way ANOVA with between-subjects factor GROUP (HIIT, MIIT, LIIT, REST) was conducted on the *first block* of visuomotor rotation task practice following AEX/Rest.

## RESULTS

### Aerobic exercise/rest data

#### Incremental exercise test and IPAQ data

Incremental exercise test results are displayed in Table 1, with complementary information in supplementary material Figure 1 and Table 1. Participants demonstrated a large range of physical activity levels, as there was a group average METS score of 6631 ± 6730 for all participants. There was no significant difference in peak power output achieved during the incremental test showed (*F_3,76_* = 1.89, *p* = .138, *η²_p_* = .07) nor physical activity levels (*F_3,76_* = 0.49, *p* = .687, *η²_p_* = 0.02) between groups.

#### Acute aerobic exercise/rest sessions

Target AEX intensity (LIIT: 49 ± 21 W; MIIT: 95 ± 31 W; HIIT: 136 ± 32 W) and active recovery (LIIT: 36 ± 17 W; MIIT: 37 ± 20 W; HIIT: 39 ± 18 W) was employed in accordance with the incremental test results. HR and perceptual response data during exercise/rest sessions, along with results, are displayed in Supplementary Tables 1 and 2.

#### Heart rate data (Fig. 3 B, C)

There was a main effect of GROUP for raw heart rate (*F_3,76_* = 212.54, *p* < .001, *η²_p_* = .89) and heart rate data as a percentage of peak value ([%HR]; *F*_3,76_ = 155.35, *p* < .001, *η²_p_* = .86), with post hoc analyses demonstrating that the HIIT group showed the highest average HR and %HR, followed by the MIIT, LIIT, and REST groups (all *ps* < .001).

**Fig.3.**
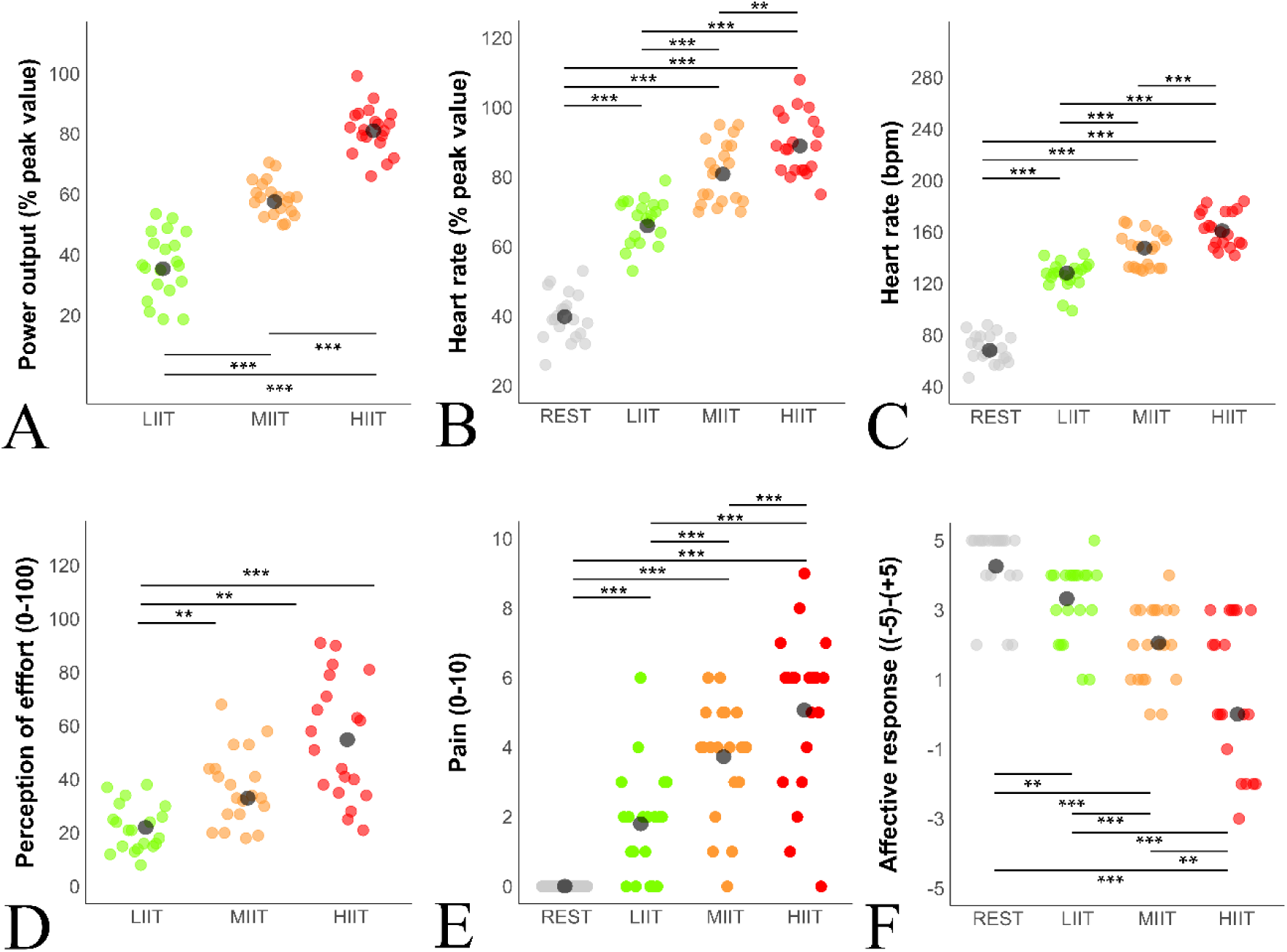
Exercise-related data: (A) Displays the power output normalized to the peak value reached during the incremental exercise test. (B) Displays the heart rate data normalized to the peak value reached during the incremental exercise test. (C) Displays the raw heart rate data in bpm. (D) Displays the perception of perceived effort data on CR-100 scale (0-100). (E) Displays the reported muscle pain data on a 0-10 scale. (F) Displays the affective response to exercise assessed using the Feeling Scale. Large dark grey circles represent the average for each condition (REST, LIIT, MIIT and HIIT), and the colored dots represent the individual data. <u>Abbreviations</u>: % peak value: percent of the peak value; bpm: beat per minute; HIIT: high intensity interval training cycling acute exercise; LIIT: light intensity interval training cycling acute exercise; MIIT: moderate intensity interval training cycling acute exercise. ** p < .01; *** p < .001.

#### Perceptual response data (Fig. 3 D, E, F)

A main effect of GROUP was found for perception of effort (*F*_2,57_ = 17.22, *p* < .001, *η²_p_* = .38), leg muscle pain (*F*_3,76_ = 40.24, *p* < .001, *η²_p_* = .61), and affective response (*F_3,76_* = 26.29, *p* < .001, *η²_p_* = .51). Post hoc analyses indicated that the HIIT group reported the highest perception of effort and muscle pain, followed by the MIIT, LIIT and REST groups (all *p*s < .007). Regarding affective response, the HIIT group reported the lowest score, followed by the MIIT, LIIT and REST groups (all *p*s < .015).

### Visuomotor reaching task data

#### Aligned visual feedback reaching

For aligned feedback reaching (Fig. 4A, C, E; *dotted horizontal lines*), there was no main effect of GROUP for reaction time (*F_3,76_* = 2.02, *p* = .119, *η²_p_* = .07), movement time (*F_3,76_* = 0.10, *p* = .959, *η²_p_* = .00), or angle at peak velocity (*F_3,76_* = 0.06, *p* = .981, *η²_p_* < .01).

**Fig.4.**
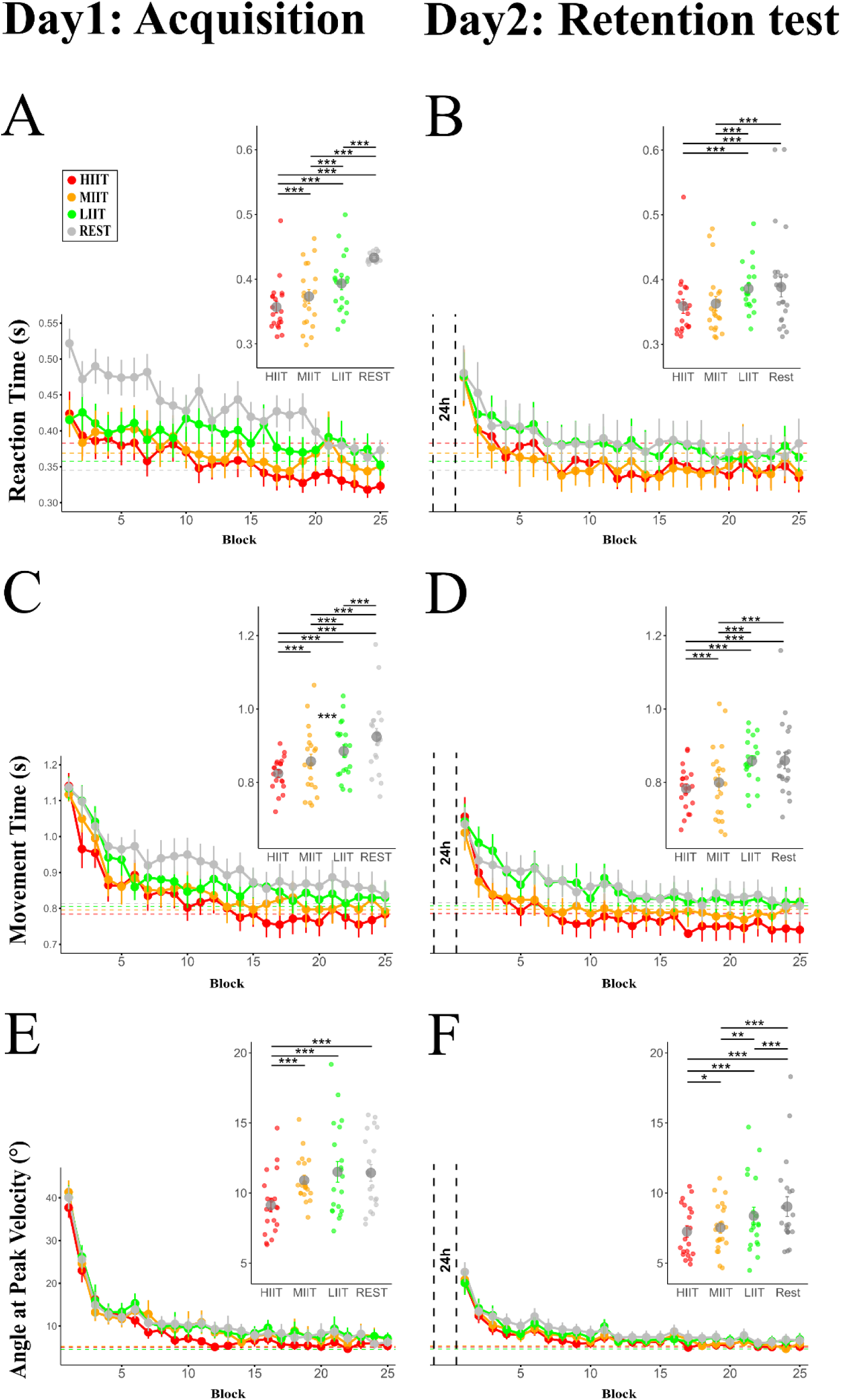
Visuomotor rotation data: *All data are reported as Mean ± standard error (SE).* **(A, B)** Display average reaction time in seconds at acquisition **(A)** and retention test **(B)** through blocks (**A, left panel; B, left panel**) and for each group (**A, right panel; B, right panel**). Horizontal lines show significant differences between the groups represented by their color, with grey for Rest, green for LIIT, orange for MIIT, and red for HIIT. **(C, D)** Display average movement time in seconds at acquisition **(C)** and retention test **(D)** through blocks (**C, left panel; D, left panel**) and for each group (**C, right panel; D, right panel**). **(E, F)** Display the average angle at peak velocity in degrees at acquisition **(E)** and retention test **(F)** through blocks (**E, left panel; F, left panel**) and for each group (**E, right panel; F, right panel**). <u>Abbreviations</u>: ***°***: degrees; **h**: hours; **HIIT**: high intensity interval training cycling acute exercise; **LIIT**: light intensity interval training cycling acute exercise; **MIIT**: moderate intensity interval training cycling acute exercise; **s**: seconds. Between-subjects: * p < .05; ** p < .01; **\*\*\*** p < .001.

#### Visuomotor rotation task: adaptation and motor learning

Group average (Fig. 4) and individual data (Supplemental Fig. 2) for visuomotor rotation task performance are displayed and the effects sizes *(Cohen’s d*) for all related post-hoc comparisons are shown in **Table 2**.

**Table 2.**
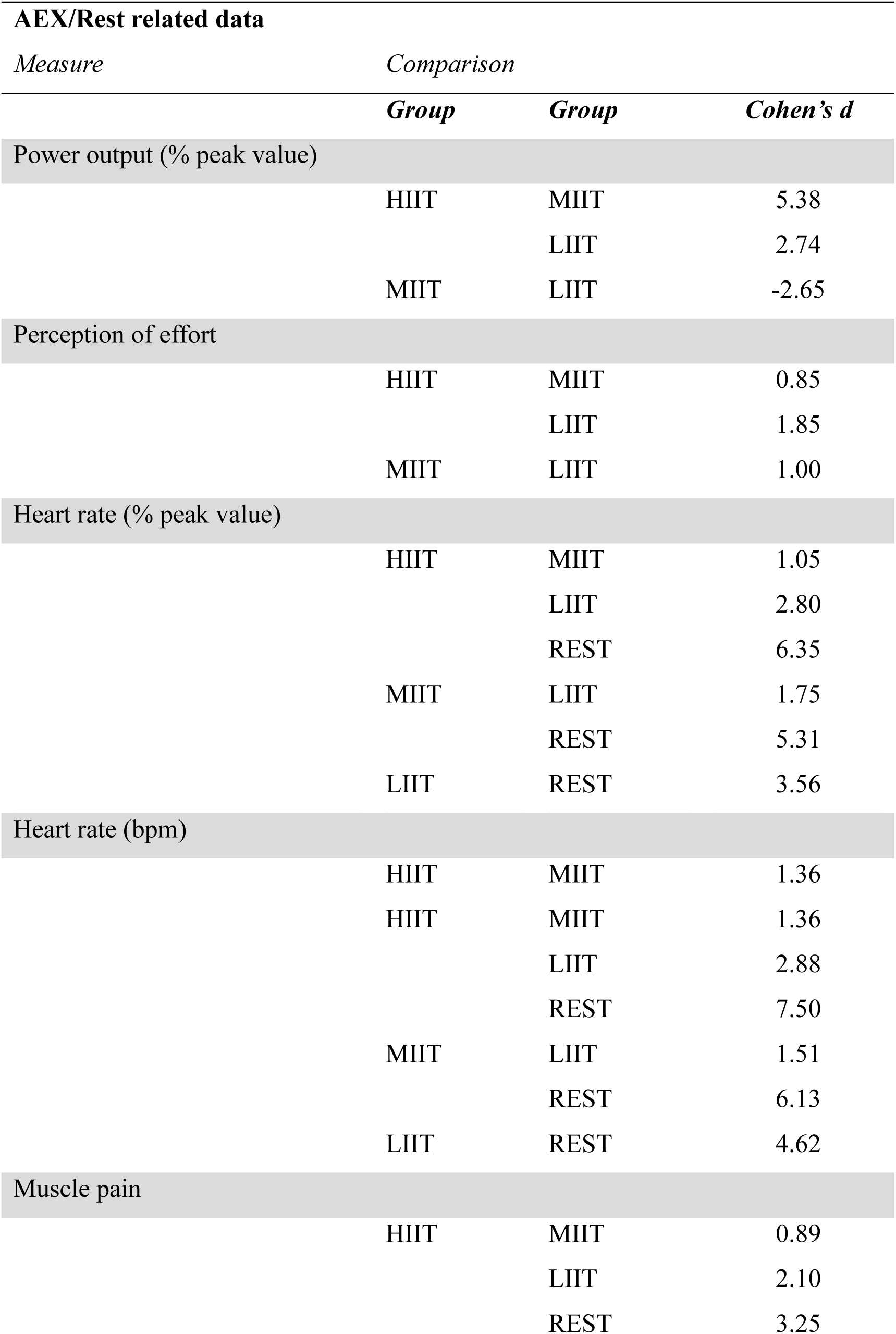
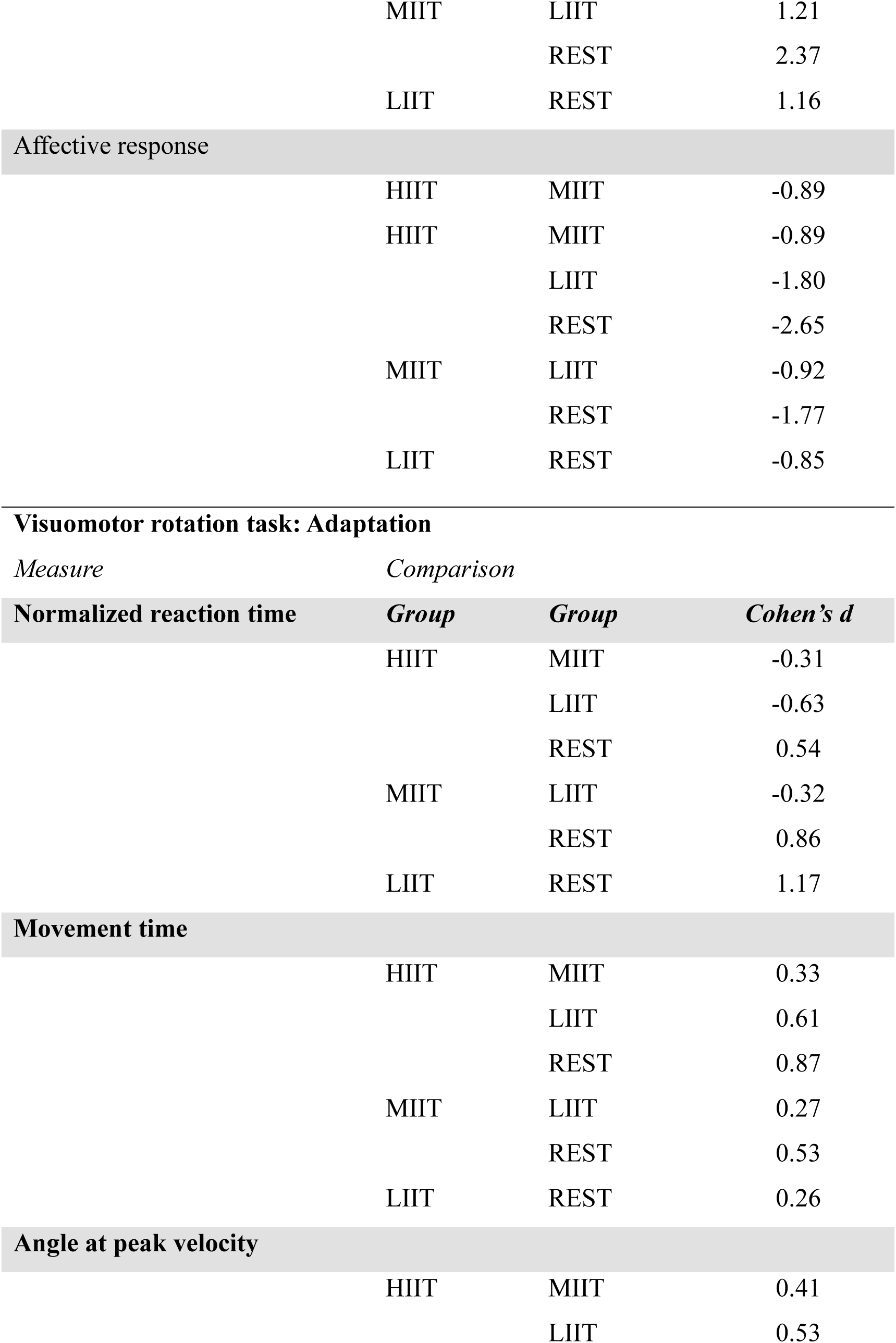
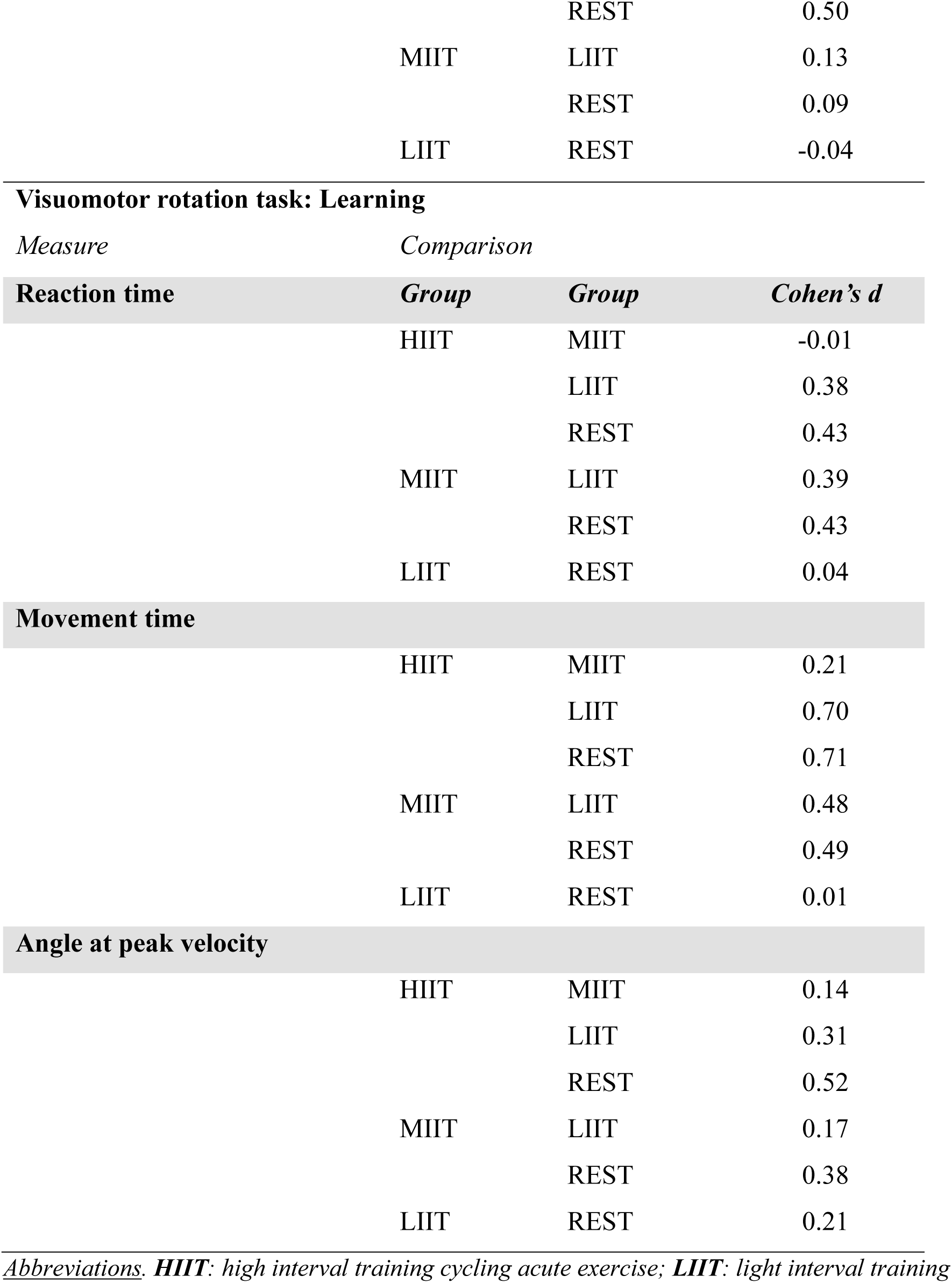
Post-hoc effect sizes (Cohen’s d) for the visuomotor rotation task data.

#### Adaptation

For reaction time, there was a GROUP × BLOCK interaction (*F_3,24_* = 1.59, *p* = .001, *η²_p_* = .06) where post-hoc analysis showed faster reaction times for all AEX groups compared to REST during the three first blocks (all *p*s < .001), and faster reaction time following HIIT compared to LIIT at block 24 (*p* < .025). Additionally, we found a main effect of GROUP (*F_3,24_* = 166.01, *p* < .001, *η²_p_* = .21) and BLOCK (*F_3,24_* = 14.92, *p* < .001, *η²_p_* = .16).

For movement time, there was a main effect of GROUP (*F_3,24_* = 68.26, *p* < .001, *η²_p_* = .10). Post-hoc analysis revealed differences among all groups (all *p*s < .001) with the HIIT group demonstrating the fastest movement time, followed by MIIT, LIIT and REST, in terms of increasing movement time. For the *first block* of visuomotor rotation task practice, there was no GROUP effect (*F_3,76_* = 0.73, *p* = .540, *η²_p_* = .03), indicating that faster movement times emerged with continued practice for the different groups. There was a main effect of BLOCK (*F_3,24_* = 41.52, *p* < .001, *η²_p_* = .35) and no GROUP × BLOCK interaction (*F_3,24_* = 68.26, *p* = .967, *η²_p_* = .03).

For angle at peak velocity, there was a main effect of GROUP (*F_3,24_* = 30.89, *p* < .001, *η²_p_* = .05). Post-hoc analysis revealed that HIIT had a lower angle at peak velocity compared to all other groups (all *p*s < .001). For the *first block* of visuomotor rotation task practice, there was no GROUP effect (*F_3, 76_* = 1.98, *p* = .132, *η²_p_* = .07), indicating that lower angle at peak velocity emerged with continued practice for HIIT. There was a main effect of BLOCK (*F_3,24_* = 229.19, *p* < .001, *η²_p_* = .75) and no GROUP × BLOCK interaction (*F_3,24_* = 0.91, *p* = .699, *η²_p_* = .03).

#### Retention test

For reaction time, there was a main effect of GROUP (*F_3,24_* = 28.05, *p* < .001, *η²_p_* = .04). Post-hoc analysis revealed faster reaction times for the HIIT and MIIT groups compared to the LIIT and REST (all *p*s < .001) groups. There was a main effect of BLOCK (*F_3,24_* = 11.97, *p* < .001, *η²_p_* = .13) and no GROUP × BLOCK interaction (*F_3,24_* = 0.30, *p* = .999, *η²_p_* = .01).

For movement time, there was a main effect of GROUP (*F_3,24_* = 62.85, *p* < .001, *η²_p_* = .09). Post-hoc analysis revealed that the HIIT group demonstrated the fastest movement time, followed by MIIT, then by LIIT and REST, in terms of increasing movement time (all *p*s < .002). There was a main effect of BLOCK (*F_3,24_* = 19.57, *p* < .001, *η²_p_* = .20) and no GROUP × BLOCK interaction (*F_3,24_* = 0.54, *p* = .999, *η²_p_* = .02).

For angle at peak velocity, there was a main effect of GROUP (*F_3,24_* = 25.85, *p* < .001, *η²_p_* = .04). Post-hoc analysis revealed differences among all groups (all *p*s < .032), with the lowest angle at peak velocity for HIIT, followed by MIIT, LIIT, and REST, in terms of increasing angle at peak velocity. There was a main effect of BLOCK (*F_3,24_* = 68.30, *p* < .001, *η²_p_* = .46) and no GROUP × BLOCK interaction (*F_3,24_* = 0.55, *p* = .999, *η²_p_* = .02).

## DISCUSSION

This study is the first to investigate the impact of AEX across the full physiological continuum of intensity on motor adaptation and learning. We demonstrated that all AEX intensities enhanced visuomotor adaptation and learning compared to rest. Critically, our findings reveal a clear dose-response relationship between AEX intensity and motor adaptation and learning: HIIT led to the most substantial improvements across all performance measures, followed sequentially by MIIT, then LIIT. This gradient of improvement was evident during *adaptation* and the *retention test*, with higher intensities yielding more comprehensive benefits, particularly for movement accuracy. Furthermore, distinct physiological and perceptual responses exhibited a dose-response relationship with AEX intensity, confirming that we successfully manipulated exercise intensity. Our results emphasize that AEX intensity is a critical parameter for enhancing motor adaptation and learning, with potential implications in sport-related and clinical contexts. Here, we discuss the potential neural and behavioral mechanisms that may underpin our findings.

### Aerobic Exercise Enhances Motor Adaptation

A novel finding in this study was a clear dose-response relationship, with higher AEX intensities leading to progressively greater improvements in adaptation. Specifically, HIIT yielded the most substantial enhancements, uniquely decreasing angle at peak velocity (a key measure of movement accuracy), in addition to increasing movement initiation and speed. While MIIT and LIIT also enhanced adaptation, primarily through increases in movement initiation and speed, their effects were to a lesser extent than HIIT. This study extends previous work^5,17^ with a systematic investigation across a full physiological continuum of intensities (light, moderate, and high), uniquely demonstrating that all AEX intensities promote adaptation relative to rest, and importantly, that the magnitude of this enhancement is dose dependent.

The observed decreases in reaction and movement time during visuomotor adaptation following all AEX intensities, compared to rest, may be indicative of a general enhancement of either arousal or cognitive strategy^53–58^. Increased arousal can improve attentional focus and processing speed, critical for rapid motor responses and the initial stages of visuomotor adaptation^59–62^. Alternatively, AEX may have facilitated the adoption or refinement of cognitive strategies necessary to quickly adjust to the rotated visual feedback, such as explicit re-aiming strategies, which have been particularly associated with reaction time^5,63–65^. While moderate-intensity AEX has long been associated with cognitive enhancements in motor task performance^66–69^, recent evidence also highlights the capacity of higher intensity AEX to improve cognitive-related performance^54,56,70,71^. This is supported by work showing that higher intensity AEX enhances task-related activation in cognition-related areas to a greater extent than moderate and lower intensity AEX^70,72^. Further, moderate-to-high intensity AEX can enhance functional connectivity in working memory networks (e.g., prefrontal-parietal cortices, hippocampus)^73–75^, which are critical for early adaptation to visuomotor rotation tasks^76,77^. The consistent decrease in reaction time and movement time, while maintaining movement accuracy, across all AEX intensities suggests that exercise enhanced efficiency in movement planning and cognitive strategy. However, while our study extended previous work^5^ by demonstrating enhanced visuomotor rotation task performance immediately following AEX of all intensities (including MIIT and LIIT), only HIIT resulted in the combined benefit of decreased reaction and movement time along with increased movement accuracy, superior to other AEX intensities and rest. This suggests that the potential cognitive strategy or arousal-based mechanisms supporting AEX effects on visuomotor adaptation^5,23,48,77–79^ may be most pronounced with higher intensity exercise in young adults.

The dose-response effect of AEX intensity on motor adaptation, particularly the superior performance with HIIT, may be supported by underlying neurophysiological changes, which may involve motor cortex excitability and cerebellar contributions^80–83^. Acute exercise is known to modulate motor cortex excitability^84^, which is associated with practice of motor adaptation tasks^36,80–82,85^. Our recent meta-analysis demonstrated that high-intensity AEX increases corticospinal excitability and decreases intracortical inhibition, whereas moderate-intensity AEX showed consistent decreases only in intracortical inhibition^84^. Our other recent work revealed a dose-response relationship between AEX intensity and corticospinal excitability increase, favoring high-intensity, alongside decreased intracortical inhibition following all AEX intensities, with the greatest reduction following moderate intensity^86^. Additionally, changes in cerebellar inhibition play a key role in the early stages of visuomotor adaptation^36^. Our previous work showed that HIIT specifically decreased cerebellar inhibition of motor cortex excitability compared to rest^12^. Taken together, modulations in motor cortical excitability and its critical connectivity with areas like the cerebellum likely contributed to our findings. Future studies should employ assessments of motor cortex excitability and cerebellar-motor cortex interactions to further elucidate these mechanisms and their dose dependent response to AEX.

### Aerobic Exercise Enhances Motor Learning

We found that all AEX intensities (HIIT, MIIT, and LIIT) significantly enhanced motor learning compared to the rest condition, along with a clear dose-response relationship in favor of HIIT, followed by MIIT, and then LIIT. This study contributes to a growing body of literature indicating AEX can enhance motor learning^3,4,10,11,13,87^, by demonstrating the impact of AEX on the unique category of motor adaptation. Our results align with a previous meta-analysis^1^, which highlighted that higher intensity AEX tends to more robustly enhance motor memory consolidation. By systematically comparing three distinct AEX intensities, our study extends previous findings and uniquely characterizes the intensity-dependent effects on visuomotor adaptation and learning. The enhancement across all AEX intensities, even LIIT, underscores the broad potential of exercise to prime the motor system for enhanced motor learning, aligning with other research showing beneficial effects across the spectrum of AEX intensities^2,6,9,10,12,19,5,6,19^.

The dose-dependent AEX-induced enhancement of motor learning may indicate the role of underlying neurobiological processes involved in memory consolidation. These mechanisms include the impact of AEX on neurotrophic factors and neurotransmitters, all contributing to plasticity within motor-related brain regions. Circulating levels of brain-derived neurotrophic factor (BDNF)^3,84,88–95^ is key candidate mechanism that may support enhanced motor skill performance and learning following AEX^88^. While higher AEX intensities have been associated with greater circulating BDNF^96^, recent work showed that peripheral BDNF is released across the full spectrum of AEX intensities (light, moderate, or high) for several hours post-exercise^97^. This widespread BDNF release may support the AEX-enhanced motor learning observed across all intensities, including LIIT. Collectively, these studies suggest that BDNF may play a significant role in AEX-enhanced motor learning, particularly at higher intensities.

The increased release or activation of dopamine and gamma-amino-butyric-acid (GABA) may have contributed to the observed AEX-enhanced motor learning. Importantly, dopamine’s D2 receptor activity is essential for AEX-induced changes in motor skill acquisition, motor cortex excitability and GABAergic inhibition^92,99^. Acute HIIT has also been shown to increase GABA concentration within the sensorimotor cortex^95^, and modulations in GABAergic inhibition are closely associated with motor skill learning^83,94,100,101^. Our recent work corroborates a dose-dependent decrease in GABAergic inhibition following AEX across intensities, with greater reductions at moderate and high intensity^84,86^. Although speculative, our findings are suggestive that activation of neurotransmitters may have played a role in the current findings of a dose-dependent AEX-induced impact on motor learning.

### Limitations

This study has some limitations. First, there was variability in physical activity levels of our participants. Prior research suggests that the impact of AEX on motor cortex excitability is more pronounced in individuals with higher physical activity levels^102,103^, which is a key mechanism associated with motor skill practice and learning^65,104–106^. Despite this, the distribution of participants across physical activity categories was relatively comparable across our groups, and the effect size for between group comparison for physical activity levels was small (*η²_p_* = 0.02, small effect size). Therefore, we anticipate that this individual variability in daily physical activity levels did not significantly impact our primary findings. However, we acknowledge a substantial difference in peak power output between groups (η²_p_ = .07, medium effect size), despite not reaching statistical significance (p = .138). In this context, we encourage future research to replicate our study using pseudorandomized group allocation to better match peak power output between groups. Second, the overall effect sizes observed for some of our primary analyses were relatively modest. This outcome is inherent to our comprehensive study design, which included four distinct groups. This deliberate approach was essential to rigorously investigate the dose-response effect of AEX intensity (along with a no-AEX control) on motor adaptation and learning. Thus, while effect sizes may appear modest in some analyses, our design adopted a pragmatic and holistic approach to address our specific research question. Finally, to isolate the dose-response effect of AEX intensity on motor learning, we intentionally controlled exercise structure (interval training) and duration (20 minutes) across AEX groups. This methodological decision inevitably led to differences in total work performed. Consequently, it becomes challenging to disentangle the unique contributions of AEX intensity from the total work performed on motor learning outcomes. Future research could address this by designing studies that equalize total work across varying exercise intensities.

### Conclusion

This study uncovered a clear dose-response relationship between AEX intensity and motor adaptation and learning. We demonstrated that all AEX intensities consistently enhanced performance compared to rest, with HIIT yielding the most pronounced improvements, followed by MIIT and LIIT. These findings collectively emphasize that the magnitude of AEX-induced enhancement is associated with exercise intensity, particularly when other exercise parameters like structure and duration are controlled. Our results underscore the critical importance of considering intensity when prescribing AEX to prime motor adaptation and learning, offering a vital foundation to develop strategies to improve motor outcomes in sports and clinical contexts.

## Author CRediT statement

Nesrine Harroum: Investigation, Data Curation, Formal Analysis, Writing – Original Draft, Project Administration. Yasmine Mahrez: Investigation, Data Curation, Writing – Review & Editing. Benjamin Pageaux: Conceptualization, Methodology, Validation, Formal Analysis, Resources, Supervision, Writing – Review & Editing, Funding Acquisition. Jason L. Neva: Conceptualization, Methodology, Validation, Formal Analysis, Resources, Supervision, Writing – Review & Editing, Funding Acquisition.

## Acknowledgments

This work is supported by the Natural Sciences and Engineering Research Council of Canada (NSERC; RGPIN-2020-05263 to JLN). JLN receives support from the Chercheur Boursier Junior 1 award of the Fonds de Recherche du Québec—Santé (FRQS #313769). BP is supported by the Chercheur Boursier Junior 1 award from the Fonds de Recherche du Québec—Santé. NH received support from both the Centre de Recherche de L’Institut Universitaire de Gériatrie de Montréal (CRIUGM), the Faculty of Medicine and the Études supérieures et postdoctorales Université de Montréal.

