## Supplementary Material for "Aerobic Exercise Intensity: A Dose-Response Effect on Motor Adaptation and Learning"

*Supplemental files*

### International Physical Activity Questionnaire (IPAQ)

#### *IPAQ description*

Daily levels of physical activity over the 7 days preceding the preliminary visit were assessed using the International Physical Activity Questionnaire[1]. The IPAQ assesses physical activity across various domains including leisure time, household chores, gardening, as well as work- and transportation-related activities. Frequency and duration of engagement in each of these activities are collected within the IPAQ. These activities were classified according to the following categories: *i)* walking, *ii)* moderate-intensity activities and *iii)* high-intensity activities. Each type of activity was weighted by its energy requirements defined in Metabolic Equivalent of Task (MET): 3.3 METs for walking, 4.0 METs for moderate-intensity activities, and 8.0 METs for high-intensity activities[2]. Physical activity levels among participants were examined to characterize the sample of participants and to compare physical activity levels between groups.

[1] *Booth M. Assessment of Physical Activity: An International Perspective. Research Quarterly for Exercise and Sport. 2000/06/01 2000;71(sup2):114-120.. doi:10.1080/02701367.2000.11082794*

[2] Ainsworth, B. E., et al. (2000). "Compendium of physical activities: an update of activity codes and MET intensities." Med Sci Sports Exerc **32**(9 Suppl): S498-504.

### International Physical Activity Questionnaire (IPAQ)

***Supplemental Fig.1****: Fitness levels distribution*

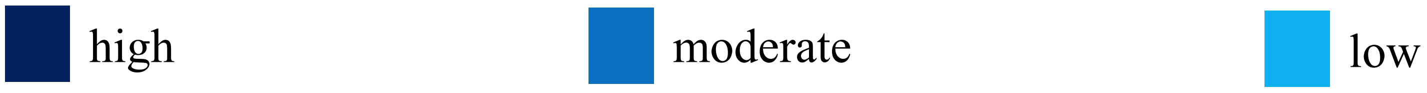


***Caption for supplemental Fig.1****:*

*This figure displays the distribution of fitness levels among groups.*

*Abbreviations: **HIIT**: high intensity interval exercise, **IPAQ**: International Physical Activity Questionnaire, **LIIT**: light intensity interval exercise, **MIIT**: moderate intensity interval exercise.*

### AEX and incremental exercise test

***Supplemental Table 1****: Results for incremental exercise test*

| | Group average |
|---|---|
| *Duration of the incremental exercise test (minutes)* | 9.66 ± 3.00 |
| *Peak power output (W)* | 155 ± 49 |
| *Heart Rate peak (bpm)* | 181 ± 2 |
| *Heart Rate rest (bpm)* | 70 ± 11 |

***Caption for supplemental Table 1****:*

*All data are reported as Mean ± SD.* ***W****: Watts,* ***bpm****: beats per minute.*

### AEX and incremental exercise test

***Supplemental Table 2****: Average power output, heart rate and perceptual responses among groups*

| ***Measure/ Group*** | **HIIT** | **MIIT** | **LIIT** | **REST** |
|---|---|---|---|---|
| *Power output (W)* | 136 ± 32 | 95 ± 31 | 49 ± 21 | NA |
| *Heart Rate (bpm)* | 162 ± 14 | 146 ± 14 | 127 ± 11 | 70 ± 11 |
| *Heart Rate (% of peak value)* | 89 ± 9 | 81 ± 9 | 67 ± 6 | 40 ± 7 |
| *Perception of effort (0 – 100)* | 53 ± 22 | 39 ± 17 | 22 ± 9 | NA |
| *Pain (0 – 10)* | 5 ± 2 | 4 ± 2 | 2 ± 2 | 0 ± 0 |
| *Affective response (-5 – (+5))* | 1 ± 2 | 2 ± 1 | 3 ± 1 | 4 ± 1 |

***Caption for supplemental Table 2****:*

*All data are reported as Mean ± SD unless otherwise noted.* ***bpm:*** *beats per minute;* ***HIIT****: high intensity interval exercise,* ***LIIT****: light intensity interval exercise,* ***MIIT****: moderate intensity interval exercise,* ***NA****: not applicable,* ***W****: Watts.*

### AEX and incremental exercise test

***Supplemental Table 3**: Statistical results for heart rate, perception of effort, muscle pain and affective responses*

| | | ***df*** | ***F*** | ***p.value*** | $\eta^2_p$ |
|---|---|---|---|---|---|
| ***IPAQ*** | Group | 3, 76 | 0.49 | 0.687 | 0.02 |
| ***Peak power output*** | Group | 3, 76 | 1.89 | 0.138 | 0.07 |
| ***Heart rate*** | Group | 3, 76 | 212.54 | **< .001** | 0.89 |
| ***Heart rate (% peak value)*** | Group | 3, 76 | 155.35 | **< .001** | 0.89 |
| ***Perception of effort*** | Group | 2, 57 | 17.22 | **< .001** | 0.38 |
| ***Muscle pain*** | Group | 3, 76 | 40.24 | **< .001** | 0.61 |
| ***Affective responses*** | Group | 3, 76 | 26.29 | **< .001** | 0.51 |

***Caption for supplemental Table 3**:*

*Abbreviations: **HIIT**: high intensity interval training session; **MIIT**: moderate intensity interval training session; **LIIT**: light intensity interval training session; **df**: degree of freedom; $\eta^2_p$: eta square partial.*

### AEX and incremental exercise test

***Supplemental Table 4**: Post hocs for heart rate, perception of effort, muscle pain and affective responses*

| | | | | *t* | *p.value* | *Cohen's d* |
|---|---|---|---|---|---|---|
| ***Heart rate (bpm)*** | HIIT | - | MIIT | 4.31 | < **.001** | 1.36 |
| | HIIT | - | LIIT | 9.10 | < **.001** | 2.88 |
| | HIIT | - | REST | 23.70 | < **.001** | 7.50 |
| | MIIT | - | LIIT | 4.79 | < **.001** | 1.51 |
| | MIIT | - | REST | 19.39 | < **.001** | 6.13 |
| | LIIT | - | REST | 14.60 | < **.001** | 4.62 |
| ***Heart rate (% peak value)*** | HIIT | - | MIIT | 3.31 | < **.01** | 1.05 |
| | HIIT | - | LIIT | 8.85 | < **.001** | 2.80 |
| | HIIT | - | REST | 20.10 | < **.001** | 6.35 |
| | MIIT | - | LIIT | 5.54 | < **.001** | 1.75 |
| | MIIT | - | REST | 16.79 | < **.001** | 5.31 |
| | LIIT | - | REST | 11.24 | < **.001** | 3.56 |
| ***Perception of effort*** | HIIT | - | MIIT | 2.69 | < **.01** | 0.85 |
| | HIIT | - | LIIT | 5.86 | < **.001** | 1.85 |
| | MIIT | - | LIIT | 3.17 | < **.01** | 1.00 |
| ***Muscle pain*** | HIIT | - | MIIT | 2.80 | < **.01** | 0.89 |
| | HIIT | - | LIIT | 6.63 | < **.001** | 2.10 |
| | HIIT | - | REST | 10.28 | < **.001** | 3.25 |
| | MIIT | - | LIIT | 3.82 | < **.001** | 1.21 |
| | MIIT | - | REST | 7.48 | < **.001** | 2.37 |
| | LIIT | - | REST | 3.66 | < **.001** | 1.16 |
| ***Affective responses*** | HIIT | - | MIIT | -2.80 | < **.01** | -0.89 |
| | HIIT | - | LIIT | -5.70 | < **.001** | -1.80 |
| | HIIT | - | REST | -8.39 | < **.001** | -2.65 |
| | MIIT | - | LIIT | -2.90 | < **.01** | -0.92 |
| | MIIT | - | REST | -5.59 | < **.001** | -1.77 |
| | LIIT | - | REST | -2.69 | < **.01** | -0.85 |

***Caption for supplemental Table 4**:*

*Abbreviations: **HIIT**: high intensity interval training session; **MIIT**: moderate intensity interval training session; **LIIT**: light intensity interval training session;. **df**: degree of freedom; $\eta^2_p$: eta square partial*

***Supplemental Fig.2*** *Indivdual visuomotor rotation data for acquisition and retention test*

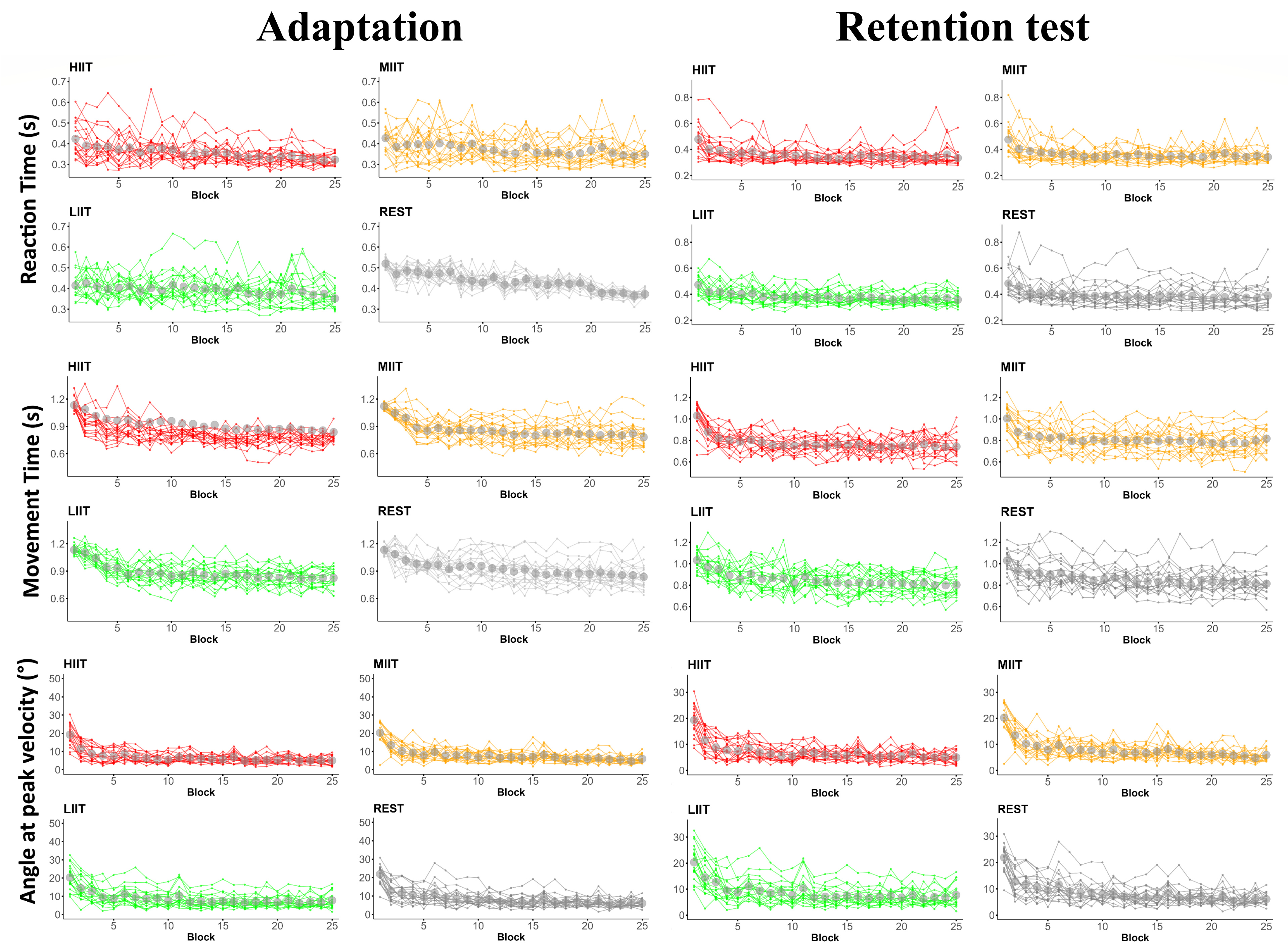


***Caption for supplemental Fig.2***:

<u>Abbreviations</u>: **°**: degrees; **HIIT**: high intensity interval training cycling acute exercise; **LIIT**: light intensity interval training cycling acute exercise; **MIIT**: moderate intensity interval training cycling acute exercise; **s**: seconds.

### No-cursor feedback reaching

#### *1. Task description*

*Participants performed reaching movements to all targets without cursor feedback for 32 trials at four time points: i) immediately before AEX/Rest, ii) after visuomotor rotation task practice, and iii) before and iv) after visuomotor rotation task practice at the 24-h retention test. In this reaching task, the white cursor representing hand movement was not visible during the reach out to peripheral targets and the reach back to the central target, each while within a 3-cm radius of the central target. This reaching task enabled the detection of sustained deviations in reaching following visuomotor rotation task practice, often referred to as aftereffects (i.e., reflecting a newly formed sensorimotor mapping)*[1].

#### *2. Data processing*

*For the no-cursor feedback reaching task, we assessed performance using angle at peak velocity to assess persistent deviation in the adapted direction without visual feedback (i.e., after effects)*[2,3].

[1] *Shadmehr R, Mussa-Ivaldi FA. Adaptive representation of dynamics during learning of a motor task. J Neurosci. May 1994;14(5 Pt 2):3208-24. doi:10.1523/jneurosci.14-05-03208.1994*

[2] *Krakauer JW. Motor Learning and Consolidation: The Case of Visuomotor Rotation. In: Sternad D, ed. Progress in Motor Control: A Multidisciplinary Perspective. Springer US; 2009:405-421.*

[3] *Kim HE, Morehead JR, Parvin DE, Moazzezi R, Ivry RB. Invariant errors reveal limitations in motor correction rather than constraints on error sensitivity. Communications Biology. 2018/03/22 2018;1(1):19. doi:10.1038/s42003-018-0021-y*

### No-cursor feedback reaching

##### *3. Statistical analysis*

*To assess the impact of AEX intensity on aftereffects via no-cursor feedback reaching, two-way mixed-model ANOVAs with between-subjects factor GROUP (HIIT, MIIT, LIIT, REST) and within-subject factor TIME (Pre, Post) were conducted on each practice session (adaptation, retention test) with angle at peak velocity as the dependent measure.*

##### *5. Results*

*Adaptation*. There was a main effect of TIME ($F_{1,3}$ = 97.92, $p$ < .001, $\eta^2_p$ = .40). Post-hoc analysis indicated greater angle at peak velocity in the adapted direction (i.e., aftereffects) post visuomotor rotation task practice compared to pre, regardless of AEX/REST group ($t_{149}$ = -9.00, $p$ < .001, $d$ = -1.58). There was no main effect of GROUP ($F_{3,3}$ = 0.77, $p$ = .515, $\eta^2_p$ = .02) nor a GROUP × TIME interaction ($F_{1,3}$ = 0.54, $p$ = .656, $\eta^2_p$ = .63).

*Retention test*. There was a main effect of TIME ($F_{3,3}$ = 249.21, < .001, $\eta^2_p$ = .03)**.** Post-hoc analysis indicated greater angle at peak velocity in the adapted direction (i.e., aftereffects) post visuomotor rotation task practice compared to pre, regardless of AEX/REST group ($t_{149}$ = -15.79, $p$ < .001, $d$ = -2.52). There was no main effect of GROUP ($F_{1,3}$ = 0.46, $p$ = .708, $\eta^2_p$ = .46) or GROUP × TIME interaction ($F_{1,3}$ = 0.43, $p$ = .729, $\eta^2_p$ = .01).

### Results for No-cursor feedback reaching

***Supplemental Table 5****: Results for the no-cursor feedback reaching*

| | | *df* | *F* | *p.value* | $\eta^2_p$ |
|---|---|---|---|---|---|
| ***Day 1 : Adaptation*** | Time | 1 | 97.92 | < .001 | 0.40 |
| | Group | 3 | 0.77 | .515 | 0.02 |
| | Time * Group | 3 | 0.54 | .656 | 0.01 |
| ***Day 2 : Retention test*** | Time | 1 | 249,21 | < .001 | 0.63 |
| | Group | 3 | 0.46 | 0.708 | 0.01 |
| | Time * Group | 3 | 0.43 | 0.729 | 0.01 |

***Caption for supplemental Table 5****:*

*For both adaptation and the retention test we only found a main effect of TIME (all ps < .001), with post hoc analyses indicating greater angle at peak velocity in the adapted direction (i.e., presence of aftereffects) post visuomotor rotation task practice compared to pre, regardless of AEX/Rest group (all ps < .001).*

### Results for No-cursor feedback reaching

***Supplemental Fig. 3***: *Results for the no-cursor feedback reaching*

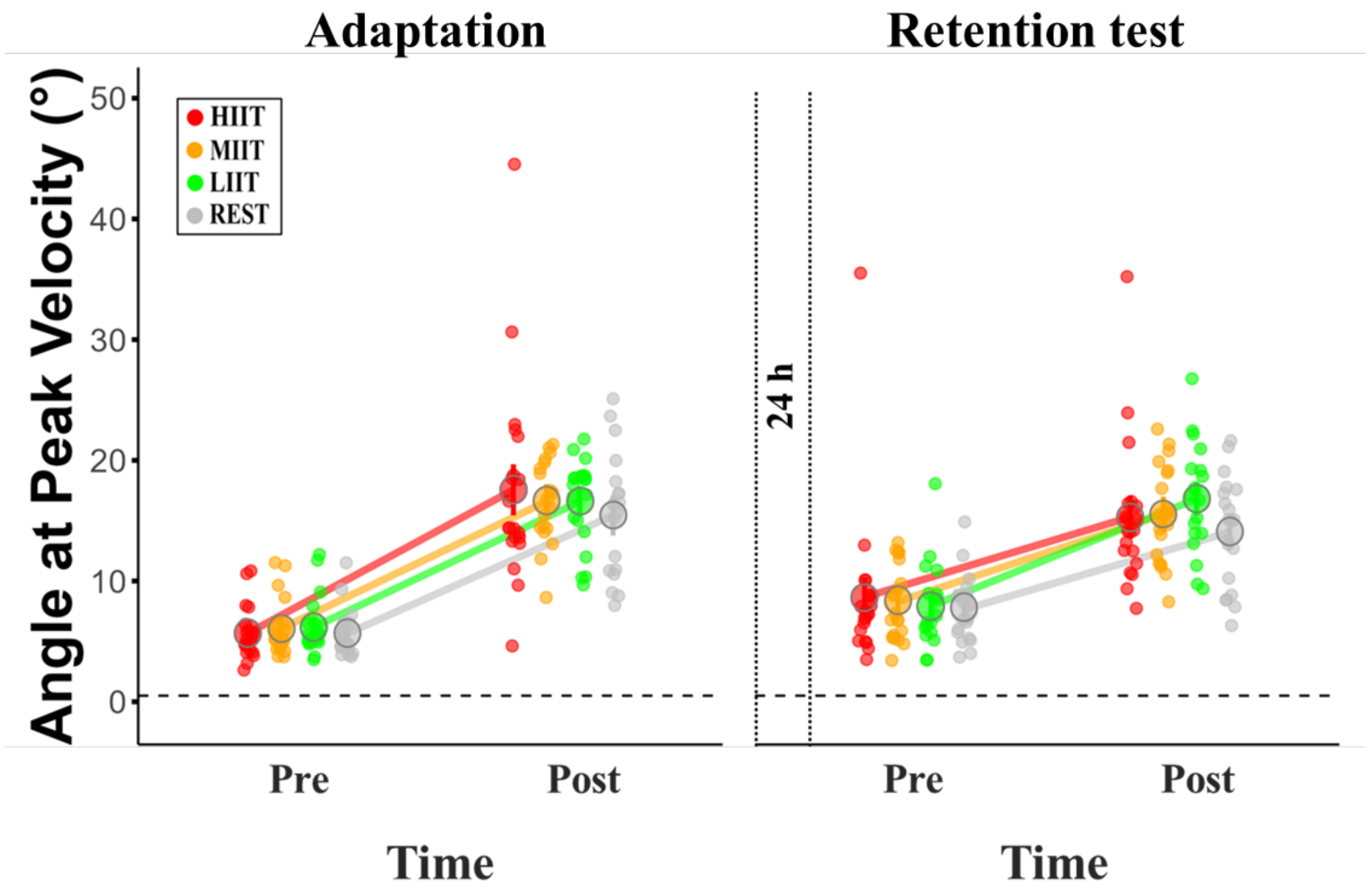


***Caption for supplemental Fig. 3***:

Average angle at peak velocity (in degrees) is shown for each group at adaptation (left panel) and retention test (right panel). Black rings represent group means. Colored dots represent individual data: red for high, orange for moderate, green for light exercise intensity, and grey for rest. For both adaptation and retention test, "Pre" and "Post" refer to measurements taken before and after the practice of the visuomotor rotation task with a 45° rotation.

Abbreviations: **°**: degrees; **h**: hours, **HIIT**: high intensity interval training cycling acute exercise; **LIIT**: light intensity interval training cycling acute exercise; **MIIT**: moderate intensity interval training cycling acute exercise; **s**: seconds.
